# Pushing the Isotopic Envelope: When carrier channels pollute their neighbors’ signals

**DOI:** 10.1101/2024.04.15.587811

**Authors:** Connor Peterson, Hannah Boekweg, Eilenora Presley, Samuel H. Payne

## Abstract

Individual cells are the foundational unit of biology, and understanding their functions and interactions is critical to advancing our understanding of health and disease. Single cell proteomics has seen intense interest from mass spectrometrists, with a goal of quantifying the proteome of single cells by adapting current techniques used in bulk samples. To date, most method optimizations research has worked towards increasing the proteome coverage of single cells. One prominent technique multiplexes many individual cells into a single data acquisition event using isobaric labels. Accompanying the single cells, one label is typically used for a mixed set of many cells, called a carrier or boost channel. Although this improves peptide identification rates, several groups have examined the impact on quantitative accuracy as more cells are included in the carrier channel, e.g. 100x or 500x. This manuscript explores how impurities in the multiplexing reagent can lead to inaccurate quantification observed as a measurable signal in the wrong channel. We discover that the severe abundance differential between carrier and single cell, combined with the reagent impurities, can overshadow several channels typically used for single cells. For carrier amounts 100x and above, this contamination can be as abundant as true signal from a single cell. Therefore, we suggest limiting the carrier channel to a minimal amount and balance the goals of identification and quantification.

## Introduction

Proteomics aims to identify and quantify proteins in biological samples. New advances in mass spectrometry and sample preparation have enabled these measurements for a single cell^1–3^. Single cell molecular measurements, including single cell proteomics (SCP), can yield insights into cellular heterogeneity and developmental/disease progression, and spatial relationships of individual cells^4^. Due to very limited ion signal, a popular method for SCP experiments is to multiplex many cells into a single data acquisition^5^. The most common commercial reagent used for multiplexing is Tandem Mass Tag (TMT)^6^, which uses a combination of ^13^C and ^15^N isotopes to create a suite of isobaric tags.

As an emerging technology, SCP is rapidly evolving to address challenges in reliability and quantitative accuracy. Using TMT-based multiplexing improves sample throughput, but has been criticized for quantitative accuracy in both bulk and single cell experiments^7–9^. The most popular experimental design, the SCoPE method^5^, uses one of the TMT channels for a “carrier,” where a large number of cells are pooled together. This dramatically increases the number of ions for MS2 spectra to boost the signal and improve peptide identification. However, the large imbalance between the abundance of the carrier and a single cell has been demonstrated to cause a number of issues with quantitative accuracy. High carrier volumes dwarf low-abundance single-cells and therefore cause poor statistical sampling and quantitative errors^10^.

A final challenge with TMT-based single cell proteomics arises from isotopic impurities in the TMT reagent, which cause a “spillover” into neighboring channels that pollutes quantitation. If a peptide is labeled with a TMT molecule that has an accidental extra ^13^C in the mass reporter region, it will cause the peptide to be counted towards quantitation of the wrong sample. The presence of impurities is well-known, and does not pose a problem with bulk proteomics. However, in SCP the carrier channel is orders of magnitude more abundant than all others and reagent impurities can have a surprisingly large effect^11^.

An unexplored question in TMT-based SCP experiments is the degree to which impurities from the carrier channel affect quantitation, specifically the possibility of spillover beyond a commonly employed single blank channel (e.g. 126 carrier paired with 127C blank). Though this would be a much smaller abundance than in 127C, the use of a very large number of cells in the carrier may be great enough to cause measurable pollution. Here, we analyze several publicly available datasets and demonstrate that this “double spillover” does occur, that the magnitude of the effect increases with increasing carrier volumes, and that it can affect the quantitative accuracy of single-cell data.

## Results

To characterize the extent to which a large carrier channel could pollute quantitation of single cell data, we identified a dataset that contained a large number of blank channels. Woo et al., introduced a new sample preparation method using nested nanowells^12^. In their experimental design, they used TMTpro 16-plex to measure protein abundance from three cell lines in triplicate. Following a common SCoPE design, label 126 was the carrier/boost channel with 10 ng or ~100 cells. Channel 127C was left empty, because this is assumed to be polluted by impurities in the 126 channel. Channel 127N was a ‘reference channel’ with 0.5 ng or ~5 cells. Channels 130N through 134N were used to label the nine samples of 0.1 ng or ~1 cell each. The remaining labels (128N, 128C, 129N, and 129C) were not used, providing an opportunity to observe the signal spillover due to reagent impurity from the carrier and/or reference.

We plotted the intensity of each TMT channel in all high-quality peptide-spectrum matches (Figure 1). Channels that were not used should not have any observable signal, unless it comes from reagent impurities. For example, a single ^13^C impurity in the 126 label will cause a measurable signal in channel 127C. This anticipated spillover is why experimental designs often leave this channel empty. As we look at Figure 1, we also notice a signal in 128C. This appears to be the result of impurity caused by the inadvertent incorporation of two ^13^C atoms into the 126 reagent. This signal is surprisingly comparable to the intensity distribution of 130N, one of the single cell samples. We similarly note that impurities in the reference channel (127N) are observed as non-negligible abundance in blank channel 128N. Using MS3 does not eliminate spillover, as the erroneous abundance is caused by the reagent impurities and not co-isolation of multiple precursors (Figure S1). The same is true for alternative placements of the carrier, e.g. the heaviest channel, as these are also found to have impurities.

**Figure 1.**
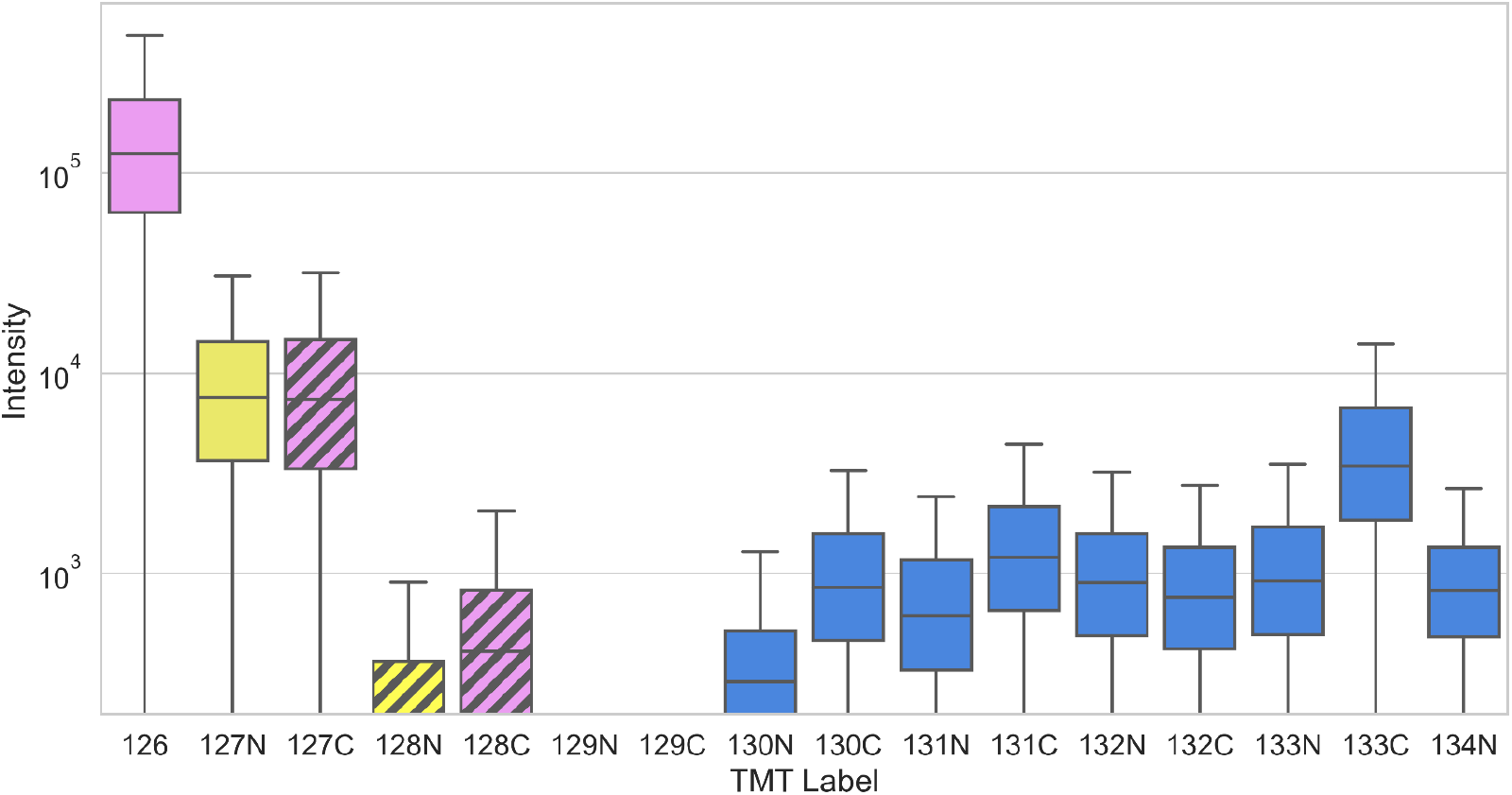
Carrier and reference samples spill over into multiple blank channels. Box plot of intensities of peptides. Channel 126 is the carrier channel (purple), with noted spillover into blank channels 127C and 128C (purple with gray striped). Channel 127N is the reference channel (yellow), with noted spillover into 128N (yellow with gray striped). All channels in blue represent single cell aliquots.

We next explored how different levels of carrier affects the spillover into 128C. Two different public datasets systematically vary the carrier channel amount relative to single cells^10,13^. These experiments do not leave the 128C channel blank, so the observed intensity is a mix of true signal from a single cell aliquot and spillover signal from impurities in the 126 reagent used for the carrier. To visualize the amount of spillover, we plot the median intensity of all PSMs for a single channel (Figure 2). These intensities are scaled to the median intensity across all channels; a value above 1 indicates that on average, intensity in this channel is more intense than other channels (see Methods). As expected, a strong correlation can be seen between the amount of carrier used and the measured abundance in channel 128C indicating the presence of spillover from reagent impurities. When carrier amounts exceed 100 times a single-cell, the reported intensities in 128C become polluted to unacceptable levels.

**Figure 2.**
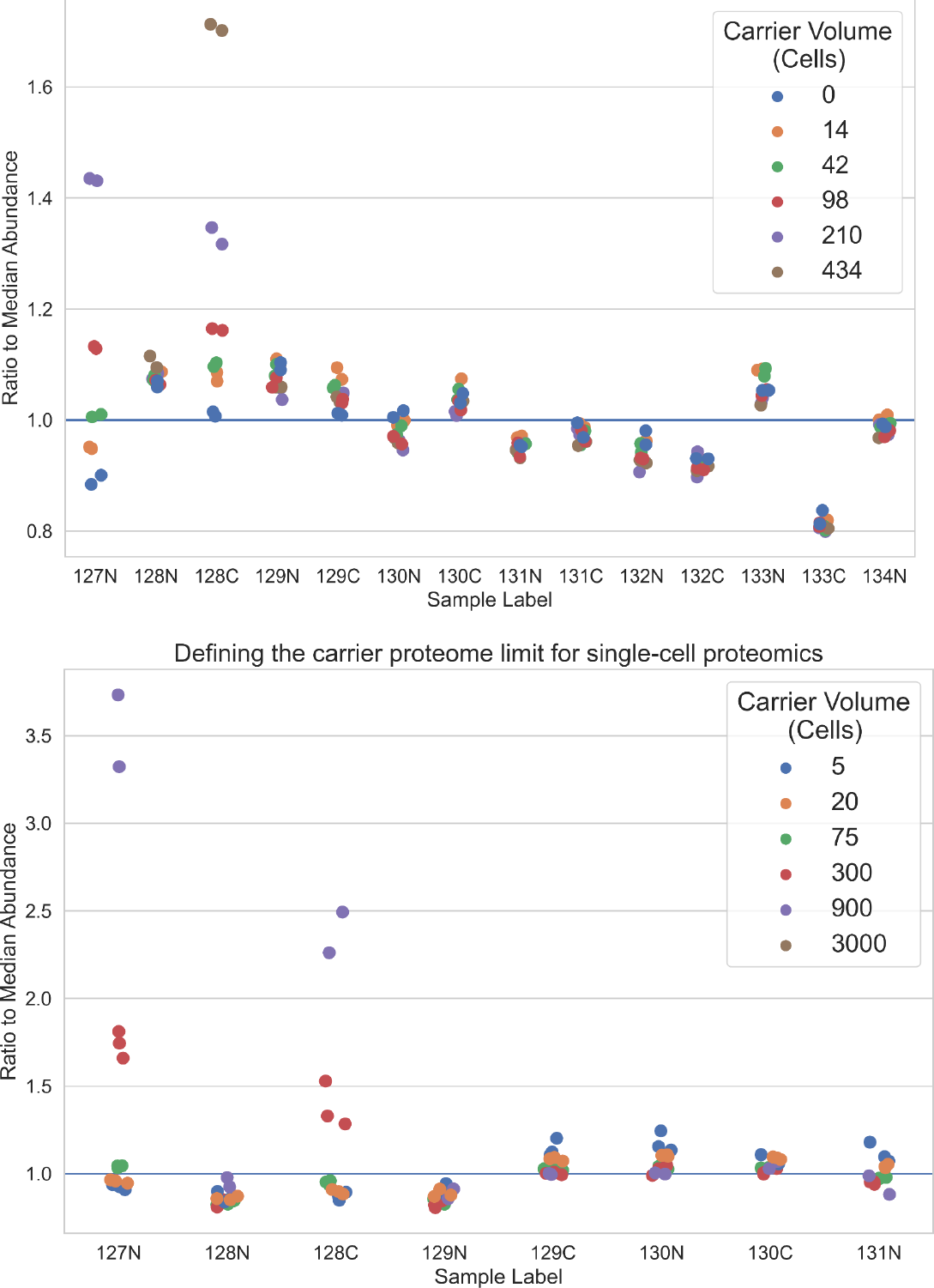
Spillover amount increases as carrier volume increases. Each plotted data point is the median of all intensities for a channel across a dataset, scaled to the median of median intensity across all channels (see Methods). The carrier channel 126 and the blank channel 127C are not graphed, as their intensity is at least an order of magnitude greater than all other channels. Data is shown for two different manuscripts: (A) for PXD027742, (B) for MSV000085959. As the volume of carrier used increases, the measured intensity in channel 128C increases accordingly. In both graphs, some datapoints are omitted, as they exist far above the plotted y-axis: channel 127C is not shown in either panel, in (A) the brown datapoints for channel 127N are not shown, in (B) brown datapoints are not shown for 127N and 128C.

To examine this spillover in real single-cell data, we examined a large experiment where 104 single cells from acute myeloid leukemia cell lines were analyzed with the standard SCoPE design^14^. As mentioned previously, we expect that impurities in the TMT reagents will lead to noticeable spillover in at least channel 128C (double ^13^C impurity from the 126 carrier channel) and 128N (single ^13^C impurity from the 127N reference channel). Differences between the 128C-labeled cell and the other cells in the experiment are thus obscured by the spillover from excessive carrier, potentially tainting biological analyses (Figure 3).

**Figure 3.**
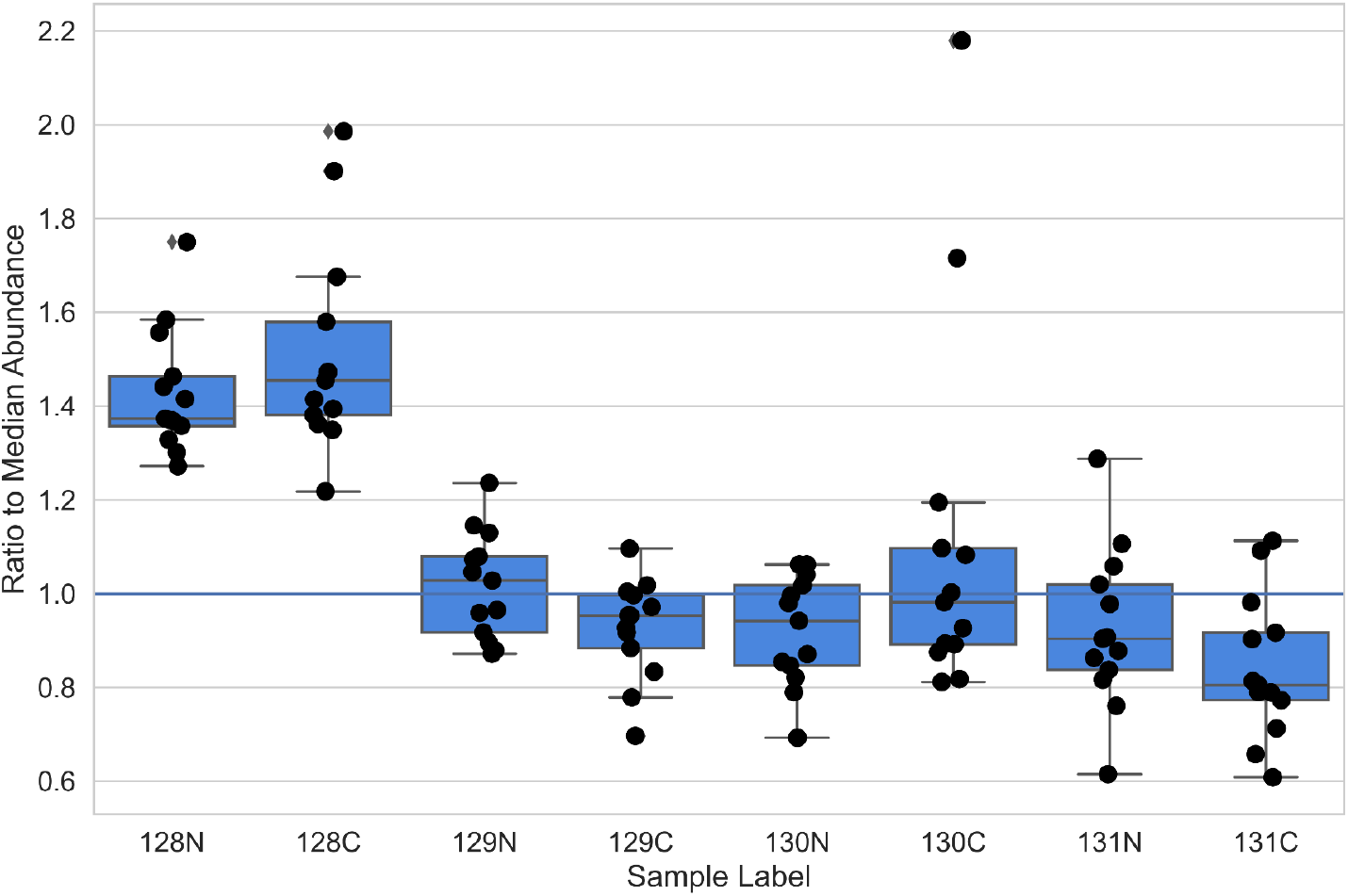
Carrier spillover affects real single-cell data. Plot showing an example of a TMT single-cell experiment using actual cells. The sample labeled with TMT128C has a significantly higher measured median abundance than other samples. Channels 126, 127C, and 127N are not shown, as their intensity far surpasses the plotted y-axis.

Channels with higher than average abundance, like those seen in Figure 3 are often corrected using a median normalization procedure. However, this assumes that the extra abundance in every PSM is a similar quantity. This assumption holds for sample loading mistakes, where one sample may accidentally have more peptides loaded. However, we were uncertain whether this is an appropriate assumption for carrier spillover. For a large carrier channel (e.g. 100x), the abundance observed in a blank channel due to spillover can be similar in magnitude to the abundance due to a single cell (see Figure 1, comparing channels 128C to 130N). This near parity in abundance is problematic. If channel 128C was a single cell and the signal due to spillover, potentially 50% of the observed intensity could be due to contamination.

For a final analysis, we wanted to characterize the variability of spillover, to help estimate the potential level of contamination in a single cell measurement. Using again the Woo et al data, we plotted the ratio of spillover intensity to carrier intensity (Figure 4). We see that for this dataset the average spillover appears to be around 6%, but is commonly as low as 4% or as high as 8%. Given the large variability in the abundance of potential spillover and its near parity in abundance compared to a single cell, we believe that median normalizations would not successfully correct for this contamination.

**Figure 4.**
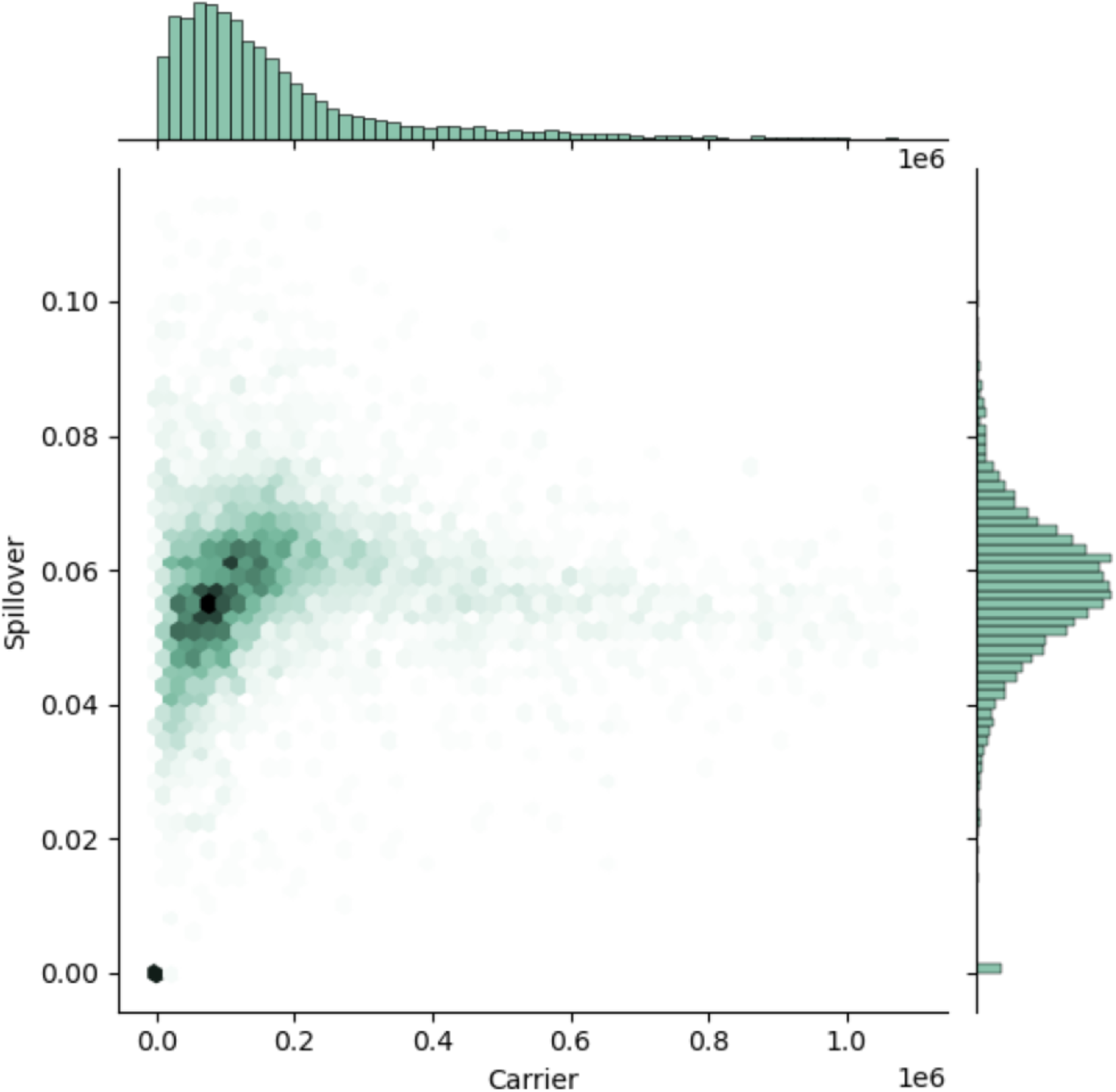
Impurity related signal spillover is variable. The amount of spillover signal, shown as the ratio between the carrier and blank channel, is plotted against the abundance of the carrier channel. No specific intensity related bias is observed, and spillover is a problem across the entire intensity range. The axis are limited to be within 3 standard deviations of the mean of their respective distributions.

## Discussion

As single cell proteomics (SCP) becomes a more common proteomics technique and gains popularity in biological inquiry, it is essential that the methodologies used to generate and analyze data are robust and accurate. Multiplexing many single cells into a single data acquisition event is a common method to meet the desired scale of single cell experiments. This method does indeed significantly increase the number of cells that can be characterized and also increase the number of quality PSMs, but has been criticized for affecting quantitative accuracy^10,13^. In this work, we identify a different way in which the use of a large carrier can compromise peptide and protein quantitation. Thus, the spillover which might contaminate single cell channels is difficult to predict; we should not rely on a general method like a median normalization to ‘correct’ for higher than expected intensity in these channels. Instead, the best option is to use fewer cells in the carrier channel and not to pollute single cell measurements.

Our results reinforce the earlier suggestions that carriers be limited to less than 100 cells to preserve quantitative accuracy. As mass spectrometry instrumentation continues to improve in ion optics and sensitivity, the carrier channel might become unnecessary.

## Acknowledgements

This work was supported by an NIGMS /NIH award (1R01GM147653) and the Simmons Center for Cancer Research.

## Methods

All code for this project is available at https://github.com/PayneLab/TMT_QC.

### Datasets and Raw Data processing

Data for this publication comes from several public single-cell proteomics datasets: Woo et al., Nat Comm 2021^12^; Ye et al., Commun Biol 2022^11^; Cheung et al., Nat Meth 2021^10^; Tsai et al., MCP 2020^14^. ProteomeXchange identifiers are noted below as used in various sections of our analysis. We note that sample metadata was not provided for these data, and so we manually created a partial metadata file for our required analyses using the sample-to-data information file (SDRF) format for each ProteomeXchange dataset.

We downloaded .raw and .mzML files of experiments of interest, and converted any .raw files to .mzML using MSConvert. We processed the .mzML files for each experiment using the MSFragger search engine via FragPipe^15^. Parameters for each analysis can be found in GitHub, ~/params/*.txt. Subsequent analyses were done using the .psm files, filtered to 1% FDR. Due to their large file size, the .psm output files cannot be stored on GitHub and are instead stored on Box (https://byu.box.com/s/dhi0wsdls0s72wt1tojz0ftmi0grc7t2). Note that our scripts used later in data processing download these files as a starting point.

### Data Analysis

The purpose of Figure 1 was to demonstrate that a large volume of carrier causes measurable signal spillover into neighboring channels. The data for Figure 1 was from Woo et al.^12^, specifically the file CellenONE_I3T_NEM_SC_Chip1_C1.raw in the MassIVE repository MSV000086809. Data output from MSFragger was plotted on a boxplot with the y-axis log10 scaled for viewing. Each column represents the measured intensities of all peptide spectrum matches (FDR < 0.01) labeled with the same TMT label. The code used to generate this figure can be found in the GitHub repository, ~/Make_Figure_1.ipynb.

Figure 2 demonstrates that the spillover into neighboring channels scales with the volume of the carrier used. Data for Figure 2a was obtained from the ProteomeXchange repository PXD027742^13^. Data used for Figure 2b was pulled from MSV000085959^10^. The .raw files that were used for each figure are listed in the corresponding SDRF file, found in our GitHub repo, ~/<proteomeXchangeID>/sdrf.xlsx. MSFragger output for all files within a dataset is combined in a single psm file. For each PSM results file, the median intensity of each TMT-labeled single-cell aliquot was calculated. The ratio of each channel’s median to the median of all channels’ medians was then calculated and then plotted with a single point, colored according to the amount of carrier utilized in that experiment. The code used to generate this figure can be found in the GitHub repository, ~/Make_Figure_2a.ipynb and ~/Make_Figure_2b.ipynb.

Figure 3 shows that the spillover problem occurs with real single-cell experiments. Data for Figure 3 was obtained from PXD017626^14^. The .raw files used in this figure are listed in its SDRF file in our GitHub repository ~/PXD017626/sdrf.xlsx. MSFragger outputs for each replicate are combined into a single psm file. The graph used for Figure 3 is similar to Figure 2. Boxplots are overlaid to more clearly illustrate the distribution of each TMT label. Code used to generate this figure can be found in our GitHub repository ~/Make_Figure_3.ipynb.

Figure 4 shows the variability in the amount of spillover, plotted against the signal intensity of the carrier channel. We used data from Woo et al.^12^, specifically the file CellenONE_I3T_NEM_SC_Chip1_C1.raw in the MassIVE repository MSV000086809. For every PSM passing 1% FDR filter, we calculated the amount of spillover as the ratio between channel 126 and 127C (carrier vs. a blank). This was plotted against the intensity of the carrier channel using a hexplot. Code used to generate this figure can be found in our GitHub repository ~/Make_Figure_4.ipynb.

## Supplemental Figures

**Figure S1.**
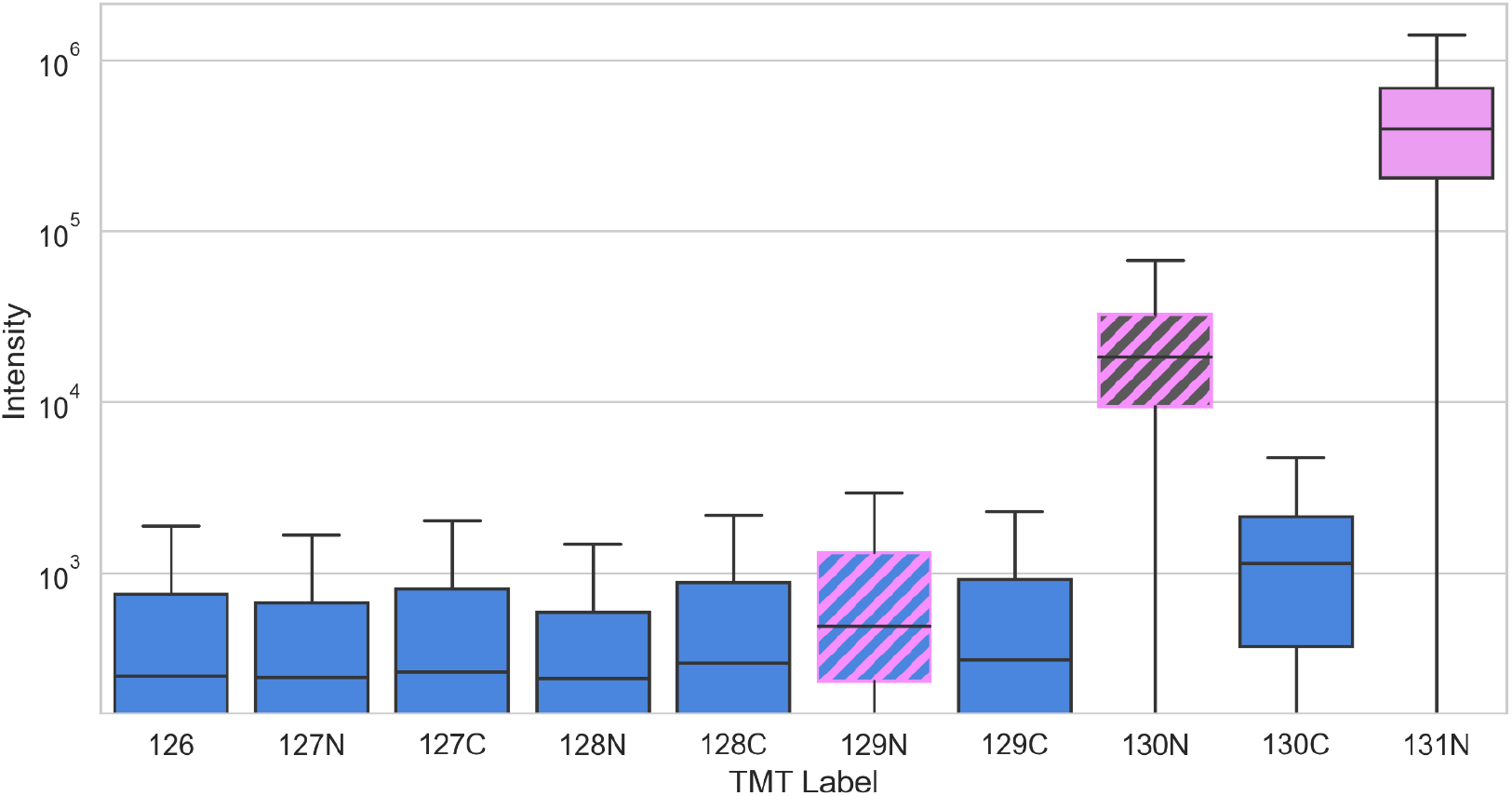
Spillover exists in MS3 data and when using the heaviest TMT label. Using data from Woo et al (see Methods), we plotted the intensity of reporter ions in each channel for all PSMs passing 1% FDR. In this plot, we only show data from MS3 spectra. The carrier channel in this experiment is 131N. Channel 130N was left blank and has significant signal due to impurities in the 131N reagent. Channel 129N was a single cell channel, but has a higher than average intensity, likely due to double impurities in the 131N reagent.

